# epDevAtlas: Mapping GABAergic cells and microglia in postnatal mouse brains

**DOI:** 10.1101/2023.11.24.568585

**Authors:** Josephine K. Liwang, Fae A. Kronman, Jennifer A. Minteer, Yuan-Ting Wu, Daniel J. Vanselow, Yoav Ben-Simon, Michael Taormina, Deniz Parmaksiz, Sharon W. Way, Hongkui Zeng, Bosiljka Tasic, Lydia Ng, Yongsoo Kim

## Abstract

During development, brain regions follow encoded growth trajectories. Compared to classical brain growth charts, high-definition growth charts could quantify regional volumetric growth and constituent cell types, improving our understanding of typical and pathological brain development. Here, we create high-resolution 3D atlases of the early postnatal mouse brain, using Allen CCFv3 anatomical labels, at postnatal days (P) 4, 6, 8, 10, 12, and 14, and determine the volumetric growth of different brain regions. We utilize 11 different cell type-specific transgenic animals to validate and refine anatomical labels. Moreover, we reveal region-specific density changes in γ-aminobutyric acid-producing (GABAergic), cortical layer-specific cell types, and microglia as key players in shaping early postnatal brain development. We find contrasting changes in GABAergic neuronal densities between cortical and striatal areas, stabilizing at P12. Moreover, somatostatin-expressing cortical interneurons undergo regionally distinct density reductions, while vasoactive intestinal peptide-expressing interneurons show no significant changes. Remarkably, microglia transition from high density in white matter tracks to gray matter at P10, and show selective density increases in sensory processing areas that correlate with the emergence of individual sensory modalities. Lastly, we create an open-access web-visualization (https://kimlab.io/brain-map/epDevAtlas) for cell-type growth charts and developmental atlases for all postnatal time points.

## Introduction

Brain growth charts provide quantitative descriptions of the brain, and are largely limited to changes in macroscopic brain volume and shape analysis during development ^1,2^. Enhanced brain growth charts that follow individual brain regions and cell types would augment our description of normal brain development as well as pathological deviations in various neurodevelopmental disorders. Brain development during the first two postnatal weeks after birth in rodents is equivalent to the late gestation through perinatal periods in humans. It features well-orchestrated and diverse events including generation and migration of neurons and non-neuronal cells, programmed cell death, and formation of synapses in a rapidly expanding brain volume ^3–6^. Specifically, γ-Aminobutyric acid-producing (GABAergic) cell development plays a critical role in establishing network excitatory-inhibitory balance in coordination with glutamatergic cells ^7–9^. For instance, cortical GABAergic neurons born in the ganglionic eminences of the embryonic brain undergo activity-dependent programmed cell death during the early postnatal period to establish expected final densities in adulthood ^10–14^. Microglia, the innate immune cells of the central nervous system (CNS), also have a critical role in brain development and wiring by facilitating GABAergic neuronal migration, developmental neuronal apoptosis, synaptogenesis, and synaptic pruning ^15–18^. Abnormal developmental processes of these cell types have been implicated in many neurodevelopmental and psychiatric disorders ^19–23^. Despite the significance of these brain cell types and emerging evidence of their regional heterogeneity ^24–26^, we have very limited information on their quantitative changes in postnatally developing brains.

Recent advances in high-resolution 3D mapping methods with cell type specific labeling make it possible to examine regionally distinct distribution of target cell types in the mouse brain ^27–31^. Previously, we discovered that GABAergic neuronal subclasses exhibit highly heterogeneous density distributions across different regions to generate distinct cortical microcircuits in the adult mouse brain ^27^. This regional heterogeneity can be partly attributed to their varying embryonic origins, birth dates, and programmed cell death. Indeed, cortical interneurons derived from both the medial (MGE) and caudal ganglionic eminence (CGE) undergo different rates of cell death ^32^. Comparably, microglia exhibit interregional and even intraregional spatial density variation across regions of the adult mouse brain, with high density in the hippocampus and basal ganglia ^33,34^. However, it remains unclear how early these region-specific density patterns emerge in the developing brain, how different GABAergic cell subclasses undergo contrasting developmental changes, and how these changes occur in synchrony with microglial development to generate the mature brain cell type landscape. One of the main challenges is the lack of developing mouse brain 3D atlases to integrate spatiotemporal trajectories of brain cell types within a consistent spatial framework ^35,36^.

Here, we create a 3D early postnatal developmental mouse brain atlas (epDevAtlas) using serial two-photon tomography (STPT) imaging at postnatal day (P) 4, 6, 8, 10, 12, and 14, along with anatomical labels based on the Allen Mouse Brain Common Coordinate Framework, Version 3.0 (Allen CCFv3) ^37^. Moreover, we develop a pipeline to systematically register target cell types in epDevAtlas and to establish standard reference growth charts for GABAergic, cortical layer-specific neuronal, and microglia cell types. Leveraging this new resource, we identified contrasting density changes of GABAergic neurons and microglia in cortical areas and white matter to gray matter colonization of microglia during early postnatal periods. Equipped with web visualization, the 3D atlas and cell type density growth charts from this study provide a suite of open science resources to understand early postnatal brain development at cellular level resolution and demonstrate the scalability of our approach to map other brain cell types.

## Results

### Creating 3D developmental brain atlases with CCFv3 anatomical labels

3D reference atlases are essential spatial frameworks that enable registration and joint analysis of different brain data ^35,37^. Here, we created morphological and intensity averaged templates using early postnatal mouse brain samples acquired by high-resolution serial two-photon tomography (STPT) imaging (**Fig. 1a**). We used Applied Normalization Tools (ANTs) to iteratively average individual samples at each age and created symmetrical templates with 20 μm-isotropic voxel size at P4, 6, 8, 10, 12, and 14 (**Fig. 1a**; see **Methods** for more details). We then applied the Allen CCFv3 anatomical labels to our new templates by performing down registration of the P56 CCFv3 template to younger brain templates using ANTs aided by manually marked major boundaries of distinct regions (e.g., midbrain-cerebellum; **Fig. 1b**; see **Methods** for more details).

**Figure 1.**
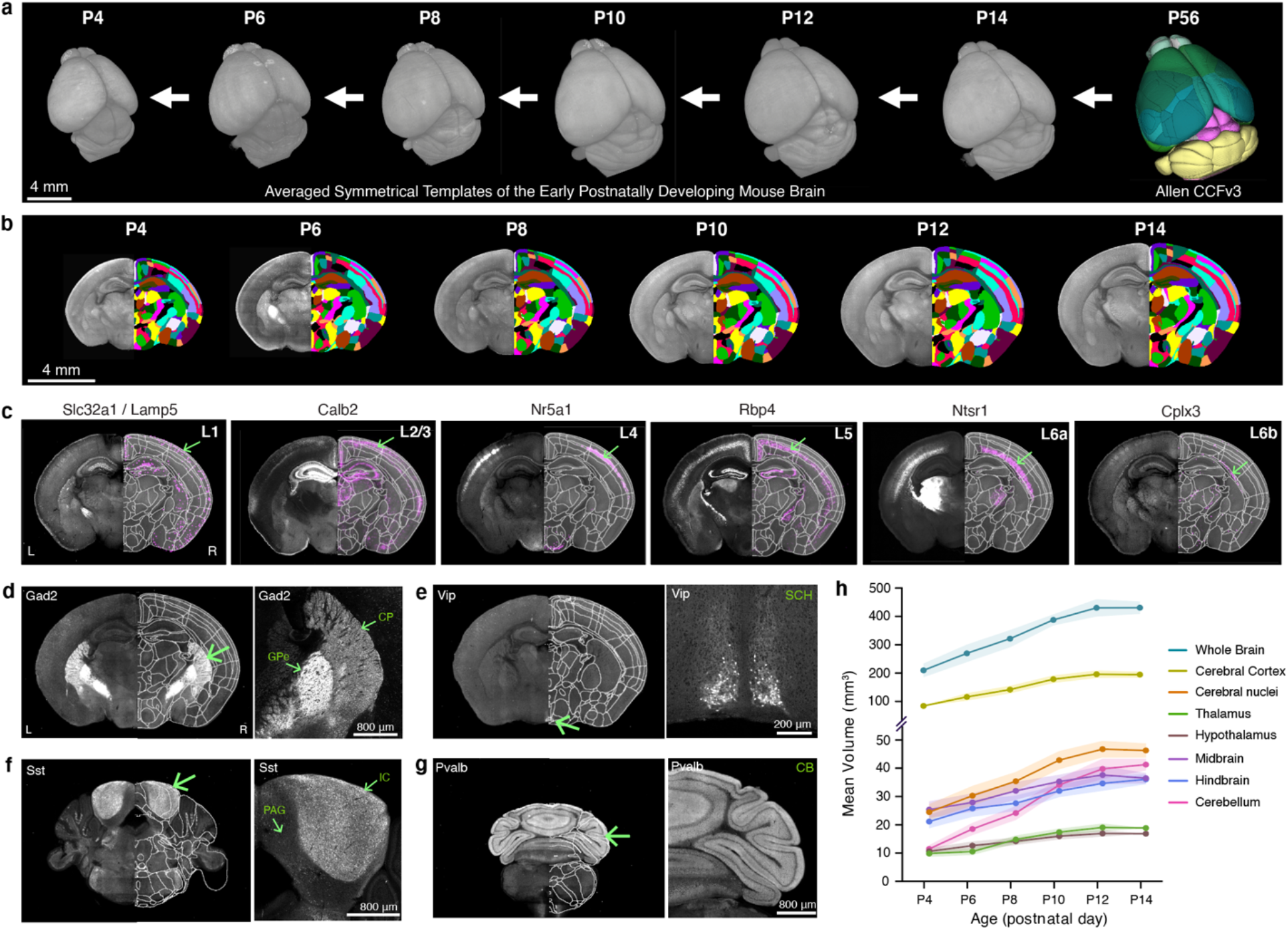
Generation of early postnatal developmental mouse brain atlas (epDevAtlas) **a-b**, Symmetrical, morphology and intensity averaged templates of the early postnatally developing mouse brain at postnatal (P) days 4, 6, 8, 10, 12, and 14 using samples from STPT imaging. Anatomical labels from the adult Allen CCFv3 were registered to each developmental time point in a stepwise manner. **c,** Cortical layer-specific cell subclass labeling using genetic strategies was implemented to refine and validate anatomical labels (L1 = layer 1, etc.). Cells are labeled by these transgenic lines by the abbreviated name of the driver lines (Table 1). Representative STPT coronal brain images shown are all from P10 animals. **d-g,** Examples of anatomical brain region delineations via cell type-specific labeling from (**d**) Gad2, (**e**) Vip, **(f)** Sst, and **(g)** Pvalb mice, collectively guiding epDevAtlas annotations. **h,** Volumetric brain growth chart during early postnatal mouse development. All brain region volumes (mm^3^) are reported as the mean ± standard deviation (s.d.; shaded area between error bars) (total n=38; see Extended Data Table 2). Additional abbreviations: CB, cerebellum; CP, caudoputamen; GPe, external globus pallidus; IC, inferior colliculus; PAG, periaqueductal gray; SCH, suprachiasmatic nucleus.

**Table 1.** List of transgenic reporter mice.

| # | Line Name | Gene | Abbreviations | Driver Mouse Line | Reporter Mouse Line | Cell Type Labeled and Considered for Study |
| --- | --- | --- | --- | --- | --- | --- |
| 1 | Gad2-IRES-Cre; Ai14 | Glutamic acid □ ecarboxylase 2 | Gad2 | Gad2-IRES-Cre | Ai14 (tdTomato) | Pan-GABAergic (Gad2-expressing) neurons |
| 2 | Sst-IRES-Cre; Ai14 | Somatostatin | Sst | Sst-IRES-Cre | Ai14 (tdTomato) | Sst-expressing neurons |
| 3 | Vip-IRES-Cre; Ai14 | Vasoactive intestinal peptide | Vip | Vip-IRES-Cre | Ai14 (tdTomato) | Vip-expressing neurons |
| 4 | Pvalb-IRES-Cre; Ai14 | Parvalbumin | Pvalb | Pvalb-IRES-Cre | Ai14 (tdTomato) | Pvalb-expressing neurons |
| 5 | Slc32a1-IRES-Cre; Lamp5-P2A-FlpO; Ai65 | solute carrier family 32 member 1/Lysosomal Associated Membrane Protein Family Member 5 | Slc32a1/Lamp5 | Slc32a1-IRES-Cre; Lamp5-P2A-FlpO | Ai65 (tdTomato) | Slc32a1/Lamp5-expressing Layer 1 cortical neurons |
| 6 | Calb2-IRES-Cre; Ai14 | Calbindin2 | Calb2 | Calb2-IRES-Cre | Ai14 (tdTomato) | Calb2-expressing Layer 2/3 cortical neurons |
| 7 | Nr5a1-Cre; Ai14 | Nuclear Receptor Subfamily 5 Group A Member 1 | Nr5a1 | Nr5a1-Cre | Ai14 (tdTomato) | Nr5a1-expressing Layer 4 cortical neurons |
| 8 | Rbp4-Cre_KL100; Ai14 | Retinol binding protein 4 | Rbp4 | Rbp4-Cre_KL100 | Ai14 (tdTomato) | Rbp4-expressing Layer 5 cortical neurons |
| 9 | Ntsr1-Cre_GN220; Ai14 | Neurotensin receptor 1 | Ntsr1 | Ntsr1-Cre_GN220 | Ai14 (tdTomato) | Ntsr1-expressing Layer 6 cortical neurons |
| 10 | Cplx3-P2A-FlpO; Ai193 | Complexin 3 | Cplx3 | Cplx3-P2A-FlpO | Ai193 (Flp: tdTomato) | Cplx3-expressing Layer 6b cortical neurons |
| 11 | Cx3cr1-GFP(+/-) | CX3C motif chemokine receptor 1 | Cx3cr1 | Cx3cr1-GFP(+/-) | - | Cx3cr1-eGFP-expressing brain microglia |

To validate and refine our registered CCFv3 labels at each age, we imaged brains from specifically selected transgenic animals. The transgenes in these animals are expressed in cell types that are differentially distributed along previously defined anatomical borders (**Table 1**) ^38,39^. We used individual or double recombinase driver lines crossed to appropriate reporter lines to label specific cell populations in the brain (**Table1; Extended Data Table 1**). For example, for cortical layers, we used a Slc32a1/Lamp5 intersectional mouse line for layer 1 (L1), Calb2 for layers 2 and 3 (L2/3), Nr5a1 for layer 4 (L4), Rbp4 for layer 5 (L5), Ntsr1 for layer 6a (L6a) and Cplx3 for layer 6b (L6b) at P6, P10, and P14 (**Fig. 1c**). Subcortical expression and axonal projections from these and additional transgenic animals also helped to validate anatomical borders for other brain regions. For instance, cortico-thalamic projections detected in Ntsr1 mouse brains specifically delineate thalamic regions (**Fig. 1c**). Moreover, Gad2 mice help to delineate substructures of the striatum, including the caudoputamen (CP) and external globus pallidus (GPe), which have markedly distinct Gad2 expression in cells and passing fibers, respectively (**Fig. 1d**). Similarly, expression patterns from Vip mice mark the suprachiasmatic nucleus (SCH) (**Fig. 1e**), while in Sst mice, the inferior colliculus (IC) is identified (**Fig. 1f**), and in Pvalb mice, the cerebellum (CB) is labeled (**Fig. 1g**). These examples, among others in multiple brain regions, helped validate, refine, and confirm the accuracy of our anatomical labels at all early postnatal ages.

Our 3D templates with ontologically consistent anatomical labels offer unique opportunities to quantify detailed volumetric changes of various brain regions at different time points (**Extended Data Table 2)**. Our data showed rapidly expanding brains with an about two-fold increase in the averaged volume of both the whole brain and the cerebral cortex between P4 and P14 (**Fig. 1h**). Moreover, the cerebellum showed the most drastic volume increase (∼four-fold) while diencephalic regions (i.e., thalamus, hypothalamus) showed the smallest increase (**Fig. 1h**). We confirm that our results closely matched previous measurements from a published MRI study ^40^, which confirms that sample preparation and STPT imaging, at this resolution, introduce insignificant volumetric changes to the mouse brain.

### Developmental mapping of GABAergic neurons

To establish cell type growth charts during the first two postnatal weeks of mouse brain development, we built a computational workflow that detects genetically defined cell types and then maps their densities on our newly developed epDevAtlas templates (**Fig. 2a**). This process involves high-resolution imaging data acquisition through STPT, machine learning (ML) cell detection based on ilastik, image registration to age-matched epDevAtlas templates, and the generation of statistical outputs detailing signals per anatomical region-of-interest (ROI) and individual ROI volumes (**Fig. 2a**) ^29^. Our cell counting pipeline produces an organized data output of volume (mm^3^), counted cells, and cell densities (cells/mm^3^) for each anatomical brain region (**Extended Data Tables 3-6**). Additionally, this workflow can be customized to work with STPT data acquired at various resolutions, as well as other high-resolution imaging data (e.g., light sheet fluorescence microscopy). Its adaptability enables the quantitative mapping of different cell types within epDevAtlas templates, which use the same anatomical labels as the widely utilized Allen CCFv3 for the adult mouse brain ^37^.

**Figure 2.**
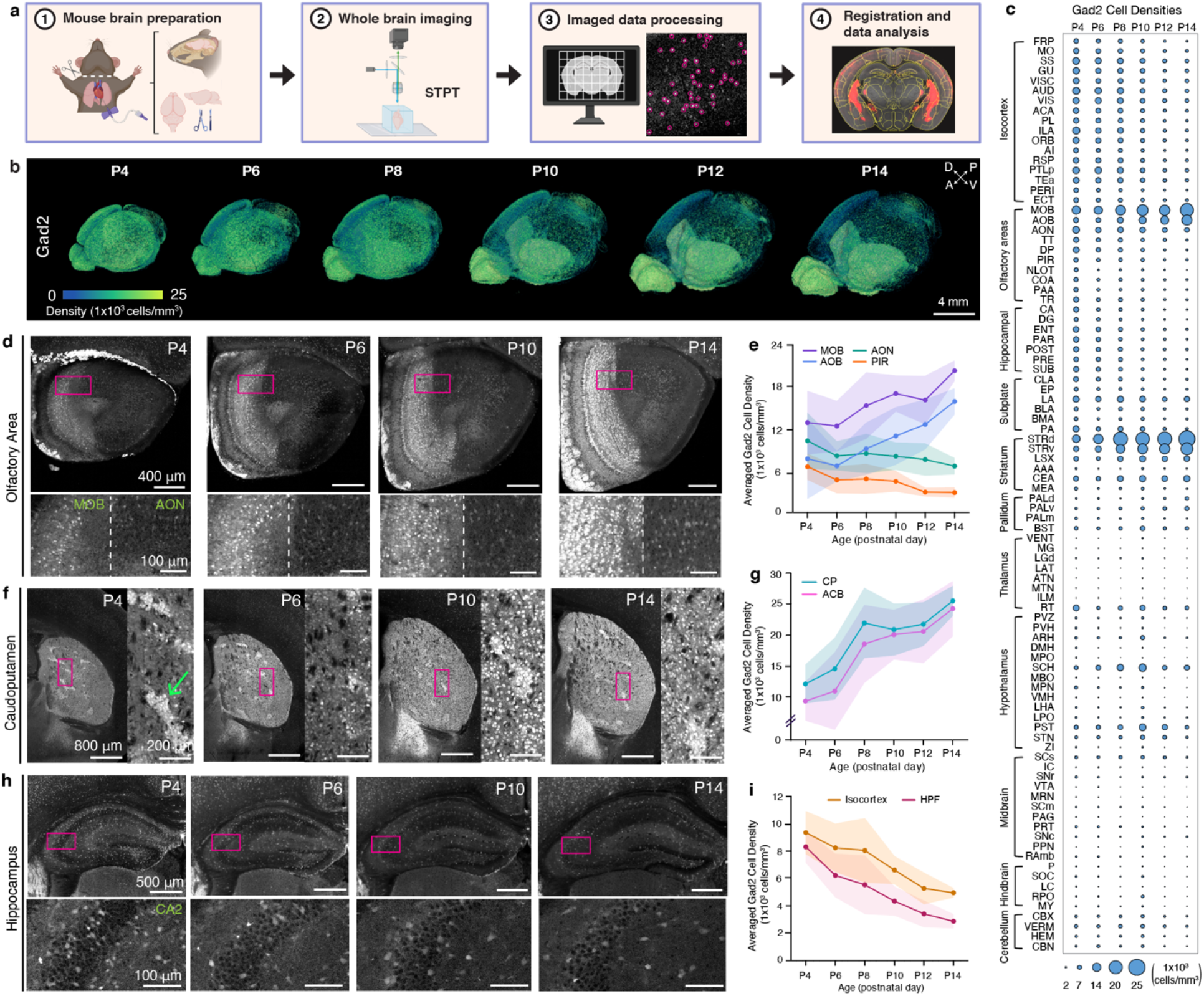
Brain-wide mapping of early postnatally developing GABAergic neurons. **a**, Overview of cell type mapping pipeline **b,** 3D renderings of Gad2 cell density (cells/mm^3^) registered to age-matched epDevAtlas. Each processed image per age is from a representative Gad2-IRES-Cre;Ai14 brain sample. **c,** Gad2 cell type growth chart across early postnatal mouse brain development, emphasizing major brain regions based on the anatomical hierarchy of Allen CCFv3. **d-e,** Olfactory areas show different trajectories of Gad2 cell density, creating a distinction between the olfactory bulb and olfactory cortices. (**d**) Representative STPT images of Gad2 cells and (**e**) their average density in the olfactory brain regions including main olfactory bulb (MOB), accessory olfactory bulb (AOB), anterior olfactory nucleus (AON), and piriform cortex (PIR) between P4 and 14. **f-g,** Striatal GABAergic cells display increased Gad2 expression from P4 to P14. (**f**) Representative STPT images of Gad2 cells and (**g**) their density in the caudoputamen (CP), with striosomes (green arrow) exhibiting earlier maturation, and nucleus accumbens (ACB). **h-i,** Hippocampal brain regions exhibit decreased Gad2 expression from P4 to P14. (**h**) Representative STPT images and (**i**) density of Gad2 cells in the hippocampal formation (HPF) and isocortex. All data in Fig. 2e, 2g, and 2i are reported as mean ± s.d. (includes shaded areas between error bars). See Extended Data Table 3 for Gad2 cell counts, density, volume measurements, and abbreviations.

As GABAergic neurons are key in setting the inhibitory tone of the brain, we first examined how this cell class changes in space and time during the first two weeks of life in mice, which is the period when genetic and external stimuli dynamically shape brain development. We applied our mapping pipeline to image the whole brain at single cell resolution in Gad2 mice and quantified the 3D distribution of the labeled cells using age-matched epDevAtlas (**Fig. 2b-c; Extended Data Table 3**). For each brain region depicted in the growth chart (**Fig. 2c**), where bubble size represents cell density, we observed temporal changes in Gad2 cell density, which fall into three main categories. We observed that from P4 to P14, Gad2 cell density either 1) continually rises, 2) declines until it levels off, or 3) remains relatively stable over time. For example, in the telencephalic area, Gad2 cell densities markedly increased in the main olfactory bulb (MOB) and the striatum (e.g., caudate putamen; CP, nucleus accumbens; ACB) (**Fig. 2d-g**), consistent with increased *Gad2* gene expression and continued neurogenesis in those areas during the early postnatal period ^41,42^. Notably, Gad2 in the CP is largely attributed to long-range projecting medium spiny neurons, which transition from the striosome at P4 to the surrounding matrix at P14 following their embryonic birth timing (**Fig. 2f**) ^43,44^. Conversely, regions such as the olfactory cortex (e.g., anterior olfactory nucleus; AON, piriform cortex; PIR) (**Fig. 2d-e**), hippocampus, and isocortex (**Fig. 2h-i**) exhibited significant reductions in Gad2 cell densities. In these cerebral cortical areas, GABAergic neurons primarily function as local interneurons. Given that subpallial striatal regions receive the main excitatory input from the cerebral cortex, the elevation of Gad2 neuronal density in the striatum and its reduction in the cortex indicate a dramatic shift in inhibitory influence in highly connected brain regions during the first two weeks of life. Gad2 cell density in other brain areas remained relatively low and stable compared to the telencephalic regions (**Fig. 2c; Extended Data Table 3**).

### Isocortical GABAergic neurons reach adult-like patterns at P12

Previous studies showed that isocortical GABAergic interneurons undergo programmed cell death in an activity-dependent manner during the early postnatal period ^10,12^. However, the timing of when regionally distinct GABAergic neuronal densities are established and when the population reaches stable adult-like spatiotemporal patterns remains unclear. Therefore, we conducted a detailed analysis of the spatiotemporal distributions of Gad2 cells in the isocortex.

In the isocortex, we observed a continuous increase in Gad2 neuronal number from P4 until P10, followed by a sharp decline until P14 (**Fig. 3a-c; Extended Data Table 3**). Throughout this period, isocortical volume showed continued growth, starting from P4, and reaching a plateau around P12 (**Fig. 3c**). We found that Gad2 cell density was highest at P4, experienced a significant decline at P10, and reached a stable level at P12 (**Fig. 3d-e**). These data are concordant with the established notion that developmentally regulated apoptosis of GABAergic cortical interneurons takes place between P1 and P15, with peak programmed cell death occurring between P7 and P11 ^10^. Previously we observed that GABAergic neurons are more densely expressed in sensory cortices compared to association areas in adult mice ^27,45^. To examine the emergence of regionally distinct Gad2 cell densities, we utilized our isocortical flatmap, which provides anatomical delineations, along with five distinct cortical domains, each represented by a different color (**Fig. 3f**) ^45^. We found that Gad2 cell density was highly enriched in sensory cortical regions (e.g., somatosensory; SS, auditory; AUD, and visual; VIS) and relatively low in association cortices (e.g., prelimbic area; PL), and this pattern was established as early as P4 (**Fig. 3d-h**). When dividing the analysis by cortical layers (L), Gad2 neurons in L2/3 exhibited the highest density at P4 and underwent the most substantial decline until P14 compared to other layers (**Extended Data Fig. 1**). Our statistical analysis showed that Gad2 level reached stable density pattern in the isocortex at P12 (**Fig. 3e**).

**Figure 3.**
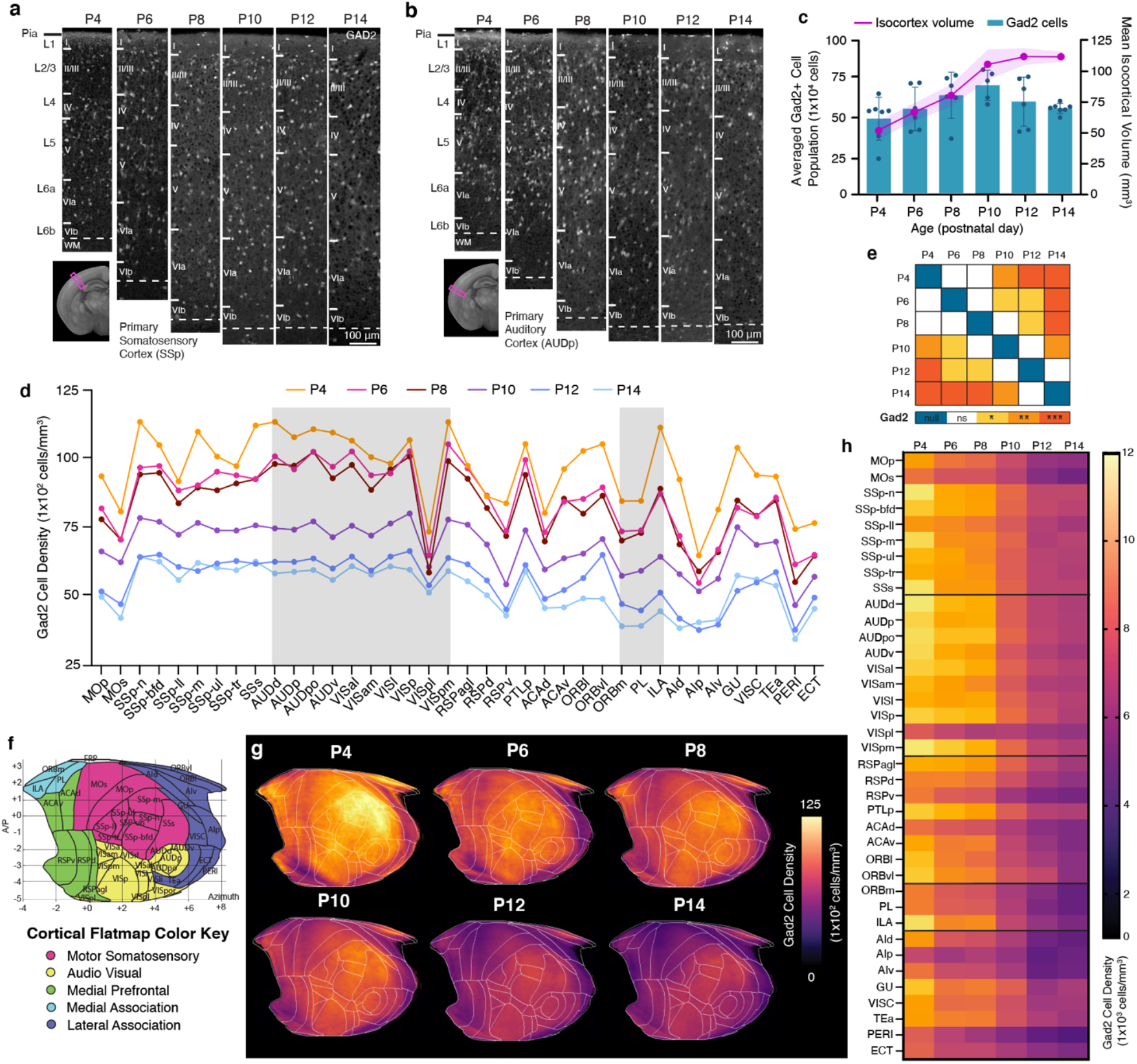
GABAergic neuronal development in the isocortex. **a-b**, Representative STPT images of Gad2 cells in the (**a**) primary somatosensory cortex (SSp) and (**b**) primary auditory cortex (AUDp). **c,** Temporal trajectory of Gad2 cell count vs. volume (mm^3^) in the isocortex. **d,** Averaged Gad2 cell density patterns across isocortical areas. Each isocortical brain region falls into one of five categories, grouped by their previously known functional and anatomical connectivity. **e,** Statistical analysis to examine significant differences between density patterns of isocortical Gad2 cells between all age pairs across (null, 0; ns, non-significant; *p < 0.05; **p < 0.005, ***p < 0.001). **f,** Isocortical flatmap with Allen CCFv3 anatomical regions and border lines. The y-axis represents the bregma’s anterior-posterior (AP) coordinates, while the x-axis indicates azimuth coordinates to combine medial-lateral and dorsal-ventral direction. **g,** Isocortical flatmaps of 3D counted and averaged Gad2 cell densities. **h,** Heatmap of averaged Gad2 cell densities. All data in Fig. 3c are reported as mean ± s.d. (includes shaded area between error bars). See Extended Data Table 3 for Gad2 cell counts, density, volume measurements, and abbreviations.

### Cortical interneurons with different developmental origins undergo differential growth patterns

Next, we began by questioning whether developmental origin can influence early postnatal density changes of cortical interneuron cell types. We assessed two specific GABAergic interneuron subclasses expressing either MGE-derived Sst or CGE-derived Vip using Sst-Cre or Vip-Cre mice crossed with Ai14 reporter mice, respectively ^9,46^.

We observed that the number of Sst interneurons steadily increased from P4 to P10 and then leveled off until P14 (**Fig. 4a-c; Extended Data Table 4**). Overall density of Sst interneurons gradually decreased over time from P4 to P14 (**Fig. 4d-e; Extended Data Table 4**). Notably, we found regionally distinct density reduction (**Fig. 4d-f**), in contrast to the relatively even reduction observed in Gad2 density across different cortical areas (**Fig. 3d**). For instance, Sst neuronal densities in medial (e.g., infralimbic: ILA) and lateral association cortices (e.g. temporal association area; TEa, perirhinal area; PERI, agranular insular area; AI) showed a dramatic reduction, where they were most enriched at P4 (**Fig. 4b, d-e**). On the other hand, the somatosensory cortex (SS) displayed the least change (**Fig. 4d-e**). When considering cortical layers, Sst neuronal densities in L5 and L6 were the highest as early as P4, and they decreased sharply in lateral association cortices, while minimal changes were observed in superficial layers (**Extended Data Fig. 2**). Statistical analysis showed that cortical Sst neurons began reaching stable density pattern at P8 (**Fig. 4f**).

**Figure 4.**
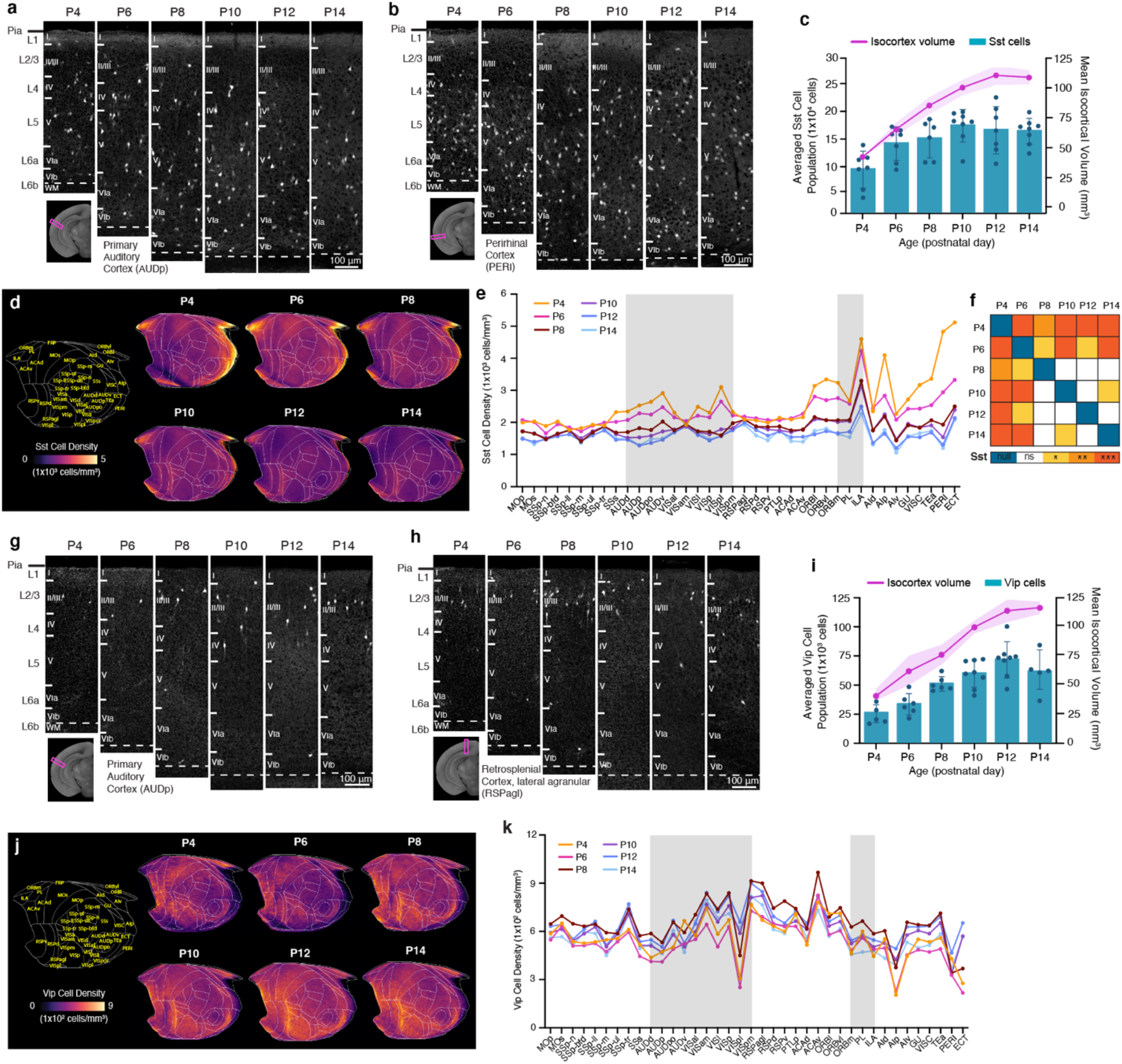
Cortical GABAergic cell types undergo differential developmental trajectories. **a-b**, Representative STPT images of Sst cells in the (**a**) primary auditory cortex (AUDp) and (**b**) perirhinal cortex (PERI). **c,** Temporal trajectory of Sst cell count vs. volume in the isocortex. **d,** Isocortical flatmaps of Sst cell densities. **e,** Averaged Sst cell density across isocortical areas. **f,** Statistical analysis to examine significant differences between density patterns of isocortical Sst cells. **g-h,** Representative STPT images of Vip cells in the (**g**) primary auditory cortex (AUDp) and **(h)** retrosplenial cortex, agranular area (RSPagl). **i,** Temporal trajectory of Vip cell count vs. volume. **j,** Isocortical flatmaps of Vip cell densities. **k,** Averaged Vip cell density (cells/mm^3^) patterns across isocortical areas. Data in Fig. 4c and 4i, are reported as mean ± s.d. (includes shaded area between error bars). See Extended Data Tables 4 and 5 for cell counts, density, volume measurements and abbreviations for Sst and Vip cells, respectively.

In contrast to Sst interneurons, CGE-derived Vip interneurons exhibited a stable cell density and regional expression patterns between P4 and P14 (**Fig. 4g-i; Extended Data Table 5**). The number of cortical Vip cells changed in tandem with cortical volume increase, resulting in no significant changes in averaged cell densities between different ages (**Fig. 4i, k**). Vip neurons are primarily enriched in the medial association (e.g., retrosplenial cortex; RSP) and audio-visual domains of the isocortical flatmap (**Fig. 4j-k**). Vip cell density was highest in L2/3 throughout the early postnatal weeks, with a relatively stable trajectory across all regions except for the lateral association cortices, in which L2/3 Vip expression was lowest at P4 and P6 before increasing until P12 (**Extended Data Fig. 2**).

These findings suggest that cortical GABAergic cell subtypes display spatiotemporal heterogeneity based on their developmental origins.

### Microglial expansion exhibits regional heterogeneity in the early postnatal mouse brain

Microglia play a pivotal role in mediating the programmed cell death of neurons and facilitating their maturation during early postnatal development ^47,48^. However, it remains unclear how microglial density evolves during early postnatal development and its connection to the early postnatal trajectory of GABAergic cells across various brain regions. To address this, we employed heterozygous Cx3cr1-eGFP(+/–) reporter mice and harnessed the epDevAtlas to systematically quantify and examine microglial distributions in the postnatally developing mouse brain.

Our comprehensive cell density mapping results unveiled significant spatial variations in the distribution of Cx3cr1 microglial populations during their early postnatal colonization of the CNS, covering the period from P4 to P14 (**Fig. 5a-b; Extended Data Table 6**). Of note, we observed the accumulation of proliferating microglia in the corpus callosum (**Fig. 5a, c**) and the cerebellar white matter (**Fig. 5a, d**). These microglial subtypes, characterized by distinct transcriptional profiles, including white matter-associated microglia (WAM) and proliferative-region-associated microglia (PAM), are situated in developing white matter tracts that regulate oligodendrocyte-mediated myelination into adulthood ^49,50^. The localized clusters of WAMs and PAMs were evident until P8 and experienced a sudden population decline by P10 (**Fig. 5a, c-d, g**). In the cerebellum, microglia are also enriched in white matter until around P10, before spreading progressively to cover other layers of the cerebellar cortex (**Fig. 5d**). Additionally, microglial morphology dynamically changes during the early postnatal period. Up until P8, microglia exhibited amoeboid morphology with thick primary branches and larger cell bodies, distinct from the more ramified microglia with comparably smaller somas observed from P10 onwards (**Extended Data Fig. 3a-b**). This morphological alteration between the first and second postnatal weeks is even more substantial in the white matter and cerebellum, implying that as white matter tracts and the cerebellar cortex mature, microglia gradually transition into a more complex and ramified phenotype (**Fig. 5d**, **Extended Data Fig. 3b**). By P12 and P14, these clonal microglia displayed a more dispersed distribution across the entire brain, forming a mosaic pattern reminiscent of tessellation (**Fig. 5a, c-f**). This mosaic-like distribution achieved by the end of the second postnatal week is then maintained into adulthood, although regional heterogeneity exists ^33,51^.

**Figure 5.**
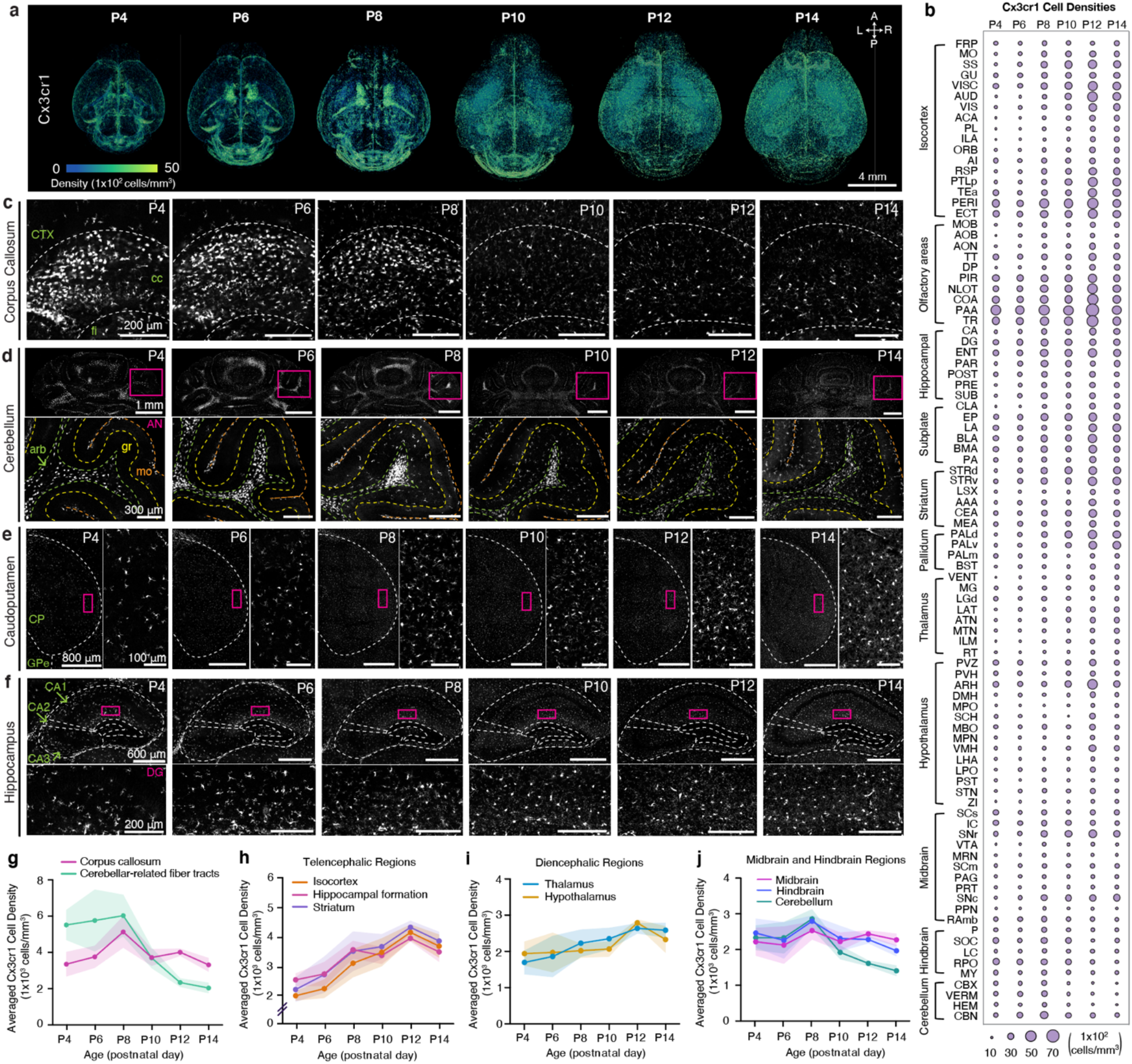
Brain-wide mapping of early postnatally developing microglia. **A**, 3D renderings of Cx3cr1 microglial cell density from representative Cx3cr1-eGFP(+/–) samples registered to age-matched epDevAtlas. **B,** Microglial (Cx3cr1) cell type growth chart **c-f,** Representative STPT images of Cx3cr1 microglia in I the corpus callosum (cc), (**d**) the cerebelluI(**e**) the caudopatamen, and (**f**) the hippocampus. Inset images in (**d-e**) are high magnification images from magenta boxed areas. **g-j,** Averaged Cx3cr1 microglial density in **(g)** white matter, specifically the corpus callosum and cerebellar-related fiber tracts, **(h)** telencephalic regions **(i)** diencephalic regions, and **(j)** midbrain and hindbrain regions. Data are reported as mean ± s.d. (shaded area between error bars). See Extended Data Table 6 for microglial cell counts, density, volume measurements, and abbreviations.

In telencephalic regions, encompassing both cortical and striatal areas, microglia displayed the most rapid expansion in density, with an approximately 200% increase from P4 to P14 (**Fig. 5e-f, h**). For example, in the striatum and hippocampus, we observed the largest increases in microglial density at the transition between the first and second postnatal weeks, with continued gradual increase until the end of the second week (**Fig. 5e-f, h**). Diencephalic regions, including the hypothalamus and thalamus, displayed modest increases in microglial density, ranging from about 50% to 100% between P4 and P14 (**Fig. 5i**). In contrast, microglial density in the midbrain and hindbrain remained relatively stable from P4 to P14, while the cerebellum showed a sharp decrease at the end of the second postnatal week, primarily due to the reduction of WAMs in the arbor vitae (arb) (**Fig. 5d, g, j**). It is also worth noting that the change in microglial morphology from amoeboid to ramified in the corpus callosum and cerebellar-related fiber tracts aligns with the drop in microglial cell density at P8 (**Fig. 5g**, **Extended Data Fig. 3**).

In summary, our results demonstrate that microglia undergo an initial expansion in selected white matter tracks, followed by further colonization of the gray matter at different rates across distinct brain regions.

### Sensory processing cortices and the dorsal striatum exhibit high microglial density

We next examined the detailed spatiotemporal patterns of microglia in the isocortex, focusing on their vital roles in fine-tuning maturing cortical inhibitory circuits ^17,18^. We discovered that the number of cortical Cx3cr1 microglia increased rapidly, surpassing the growth in isocortex volume (**Fig. 6a-c**). Therefore, the average cell density of cortical Cx3cr1 microglia exhibited a swift increase until P12 and displayed a significant negative correlation with isocortical Gad2 interneuron density (**Fig. 6d**). Our investigation also revealed regionally heterogeneous expansion of microglia (**Fig. 6e-f**). To visualize these spatial density patterns of microglia over time, we employed the isocortical flatmap (**Fig. 6g**). At P4, the flatmap highlighted the emergence of heightened Cx3cr1 density in specific focal areas within the medial and lateral association regions linked with WAMs (**Fig. 6f**). By P6, there was increased microglial density within sensory regions, particularly in the primary somatosensory cortex (SSp) and the retrosplenial cortex (RSP) (**Fig. 6e-g**). At P12 and P14, microglia began to densely populate other sensory areas, including the auditory (AUD) and visual (VIS) cortices (**Fig. 6e-g**). This creates a gradient of low-to high-density microglial distribution along the anterior-posterior axis in the isocortex by the conclusion of the second postnatal week (**Fig. 6f-g**). We observed a relatively uniform increase in microglial density across all layers from P4 to P14 (**Extended Data Fig. 4**). It is important to note that the emergence of high-density microglia in sensory cortices correlates with the onset of active sensory input, such as active whisking (linked with SSp) and the vestibular righting reflex (linked with RSP) at around P6, and the opening of ears (AUD) and eyes (VIS) at around P12 ^52–54^.

**Figure 6.**
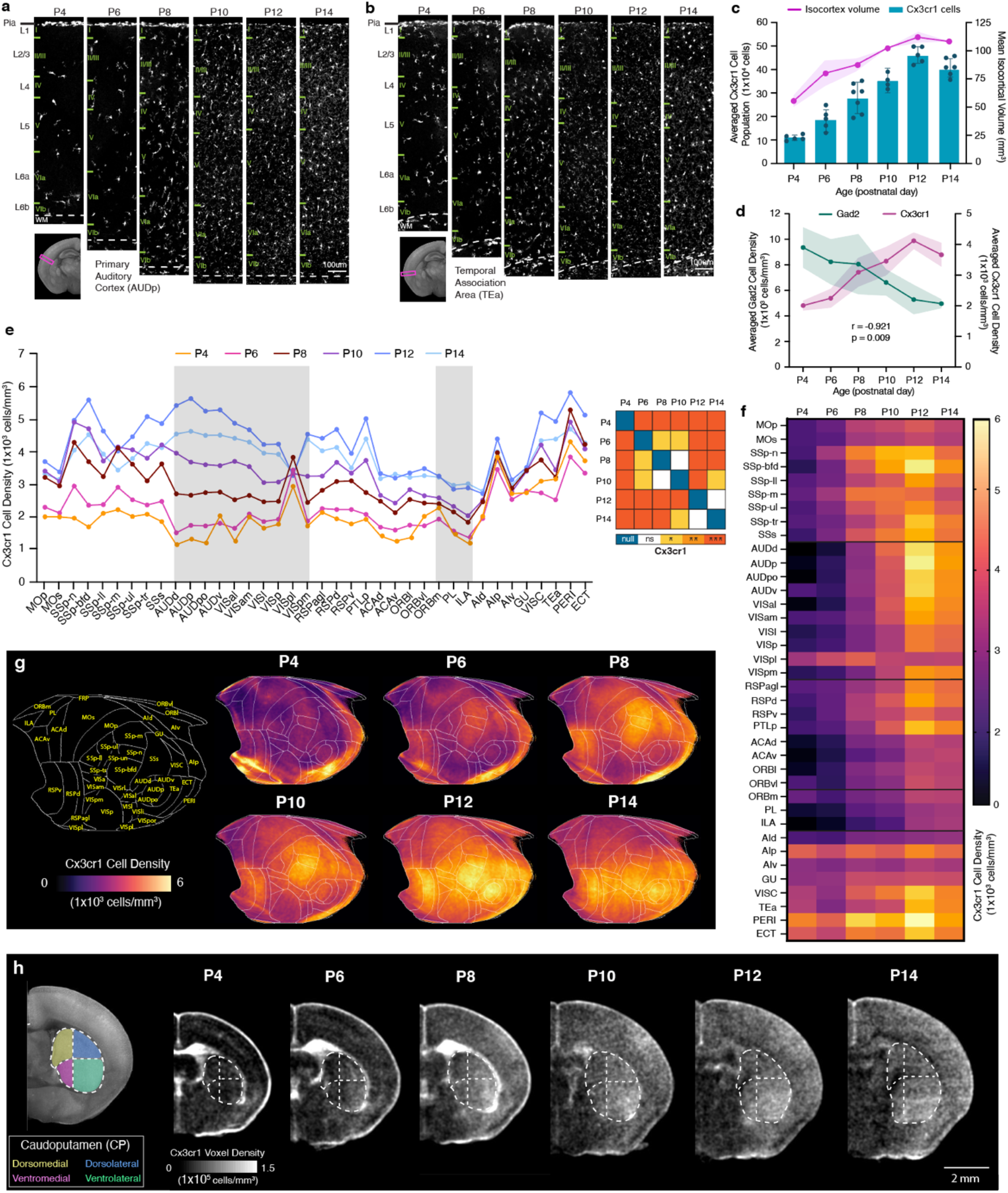
Cortical microglial expansion during early postnatal development. **a-b**, Representative STPT images of Cx3cr1 microglia in the (**a**) primary auditory cortex (AUDp) and (**b**) temporal association cortex (TEa). **c,** Temporal trajectory of Cx3cr1 microglial count vs. volume. **d,** Microglial (Cx3cr1) and GABAergic (Gad2) cell density trajectories between P4 and P14 are significantly anti-correlated. **e,** (left) Averaged Cx3cr1 microglial density patterns across isocortical areas. (right) Statistical analysis to examine significant differences between density patterns of isocortical microglia. **f,** Isocortical flatmaps of Cx3cr1 microglial densities. **g,** Heatmap of Cx3cr1 microglial densities. **h,** Striatal divisions of the caudoputamen (CP) into four functional domains (dorsomedial, yellow; dorsolateral, blue; ventromedial, magenta; ventrolateral, green) show a distinct Cx3cr1 microglial population increase in the ventrolateral CP at P4. Data in Fig. 6c-d are reported as mean ± s.d. (includes shaded area between error bars). See Extended Data Table 6 for microglial cell counts, density, volume measurements, and abbreviations.

Considering that the caudate putamen (CP), also known as the dorsal striatum, receives topological projection from distinct cortical areas ^55^, we questioned whether microglia density changes in the CP resemble developmental patterns observed in the isocortex. Indeed, we found that the ventrolateral CP, which primarily receives projections from the SSp, exhibited a higher density compared to the ventromedial CP, which receives projections from the association cortex (**Fig. 6h**)^55^. This difference in regional density was not evident at P4 and began to emerge at P6, when microglial density started to prominently populate the SSp (**Fig. 6h**).

These findings suggest that microglia preferentially increase their density in sensory processing areas as a response to activity-dependent circuit maturation, when external sensory information becomes available during the first two postnatal weeks ^56^.

### Cell type growth chart as a new resource

To enhance accessibility to epDevAtlas and its detailed cell type mappings, we developed a user-friendly web visualization based on Neuroglancer, available at https://kimlab.io/brain-map/epDevAtlas. This platform allows users to explore full-resolution images and mapped cell density data with age-matched epDevAtlas templates and labels, including cortical layer-specific reporter mice (**Fig. 7a-d**).

**Figure 7.**
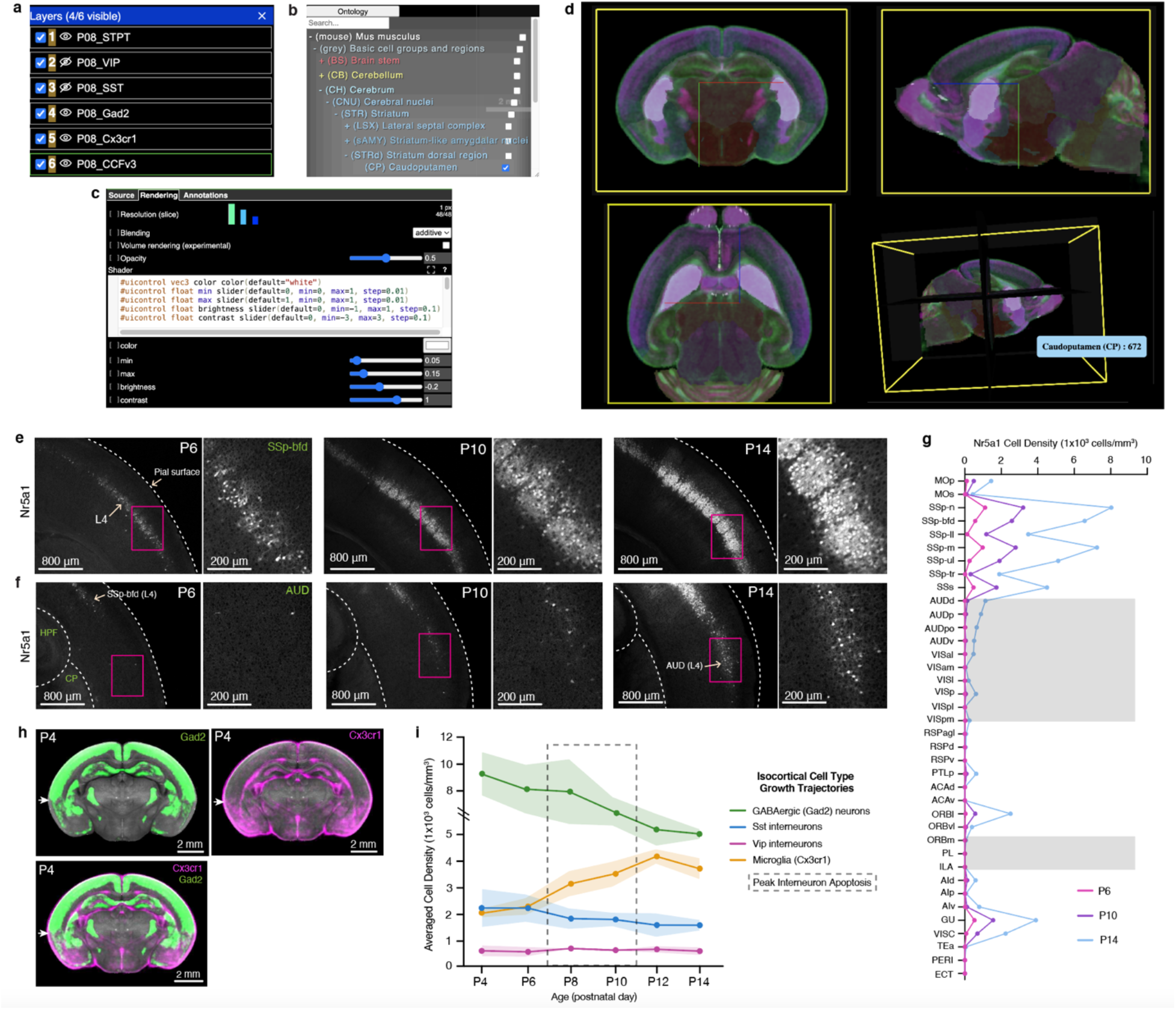
Web visualization and integrative analysis with cell type growth charts. **a-d**, Neuroglancer based visualization enables users to (**a**) select cell types or atlas/annotations, (**b**) examine anatomical label ontology, (**c**) modify visualization setting (e.g., color, intensity), (**d**) with orthogonal 3D views. **e-f,** Representative STPT images of Nr5a1 cells in layer 4 (L4) of the (**e**) primary somatosensory cortex, barrel fields, and (**f**) auditory cortex. **G,** Nr5a1 cell density across isocortical areas. **H,** Average 3D density data of Gad2 cells (top left; green; n=7) and Cx3cr1 microglia (top right; in magenta; n=5), overlaid (bottom left) registered to the P4 epDevAtlas template (gray background). White arrows highlight the border between isocortical and the olfactory cortex from the two cell types with the complementary density pattern. **€**, Summary of growth chart of GABAergic and microglial cell type densities in the isocortex.

For instance, we used Nr5a1 mice to label L4 cortical neurons. We observed an early emergence of labeled cells in the SS region of the developing cortex at P6, followed by a remarkable surge in cell density within L4 of the barrel field (SSp-bfd) at P10 that continually increased until P14 (**Fig. 7e, g**). In comparison, Nr5a1 cells within L4 of AUD and VIS regions exhibited a delay in density growth with sudden increase at P14 (**Fig. 7f, g**). This observation strongly indicates a correlation between Nr5a1 expression in cortical L4 and the developmental onset of individual sensory modalities.

Moreover, the integration of multiple cell types allows us to pinpoint regional variations in cell type compositions. Take P4 as an example, where we observed that GABAergic neurons and microglia displayed contrasting yet complementary density patterns in the isocortex and olfactory cortex (**Fig. 7h**). While GABAergic neurons maintained higher densities in the isocortex compared to the olfactory cortex, microglia exhibited the opposite pattern (**Fig. 7h**).

With this new resource, users can explore individual cell types or their combinations, facilitating comparisons of their spatial distribution across developmental stages, as summarized in the example given for the isocortex (**Fig. 7i**).

## Discussion

We present cell type growth charts of GABAergic neurons and microglia in the early postnatally developing mouse brain using the epDevAtlas as 3D STPT-based atlases. Standard biological growth charts are essential tools to comprehend normal growth and identify potential pathological deviations ^1,57^. Existing brain growth charts are largely limited to macroscopic volumetric and shape analyses. Therefore, our novel cell type growth charts significantly enhance our understanding of brain cell type composition during early development and can serve as the standard metric for evaluating alterations in pathological conditions.

The importance of 3D brain atlases as a standardized spatial framework is well recognized in integrating diverse cell census information ^58,59^. For instance, the Allen CCFv3 serves as a widely used standard adult mouse brain atlas for cell census data integration ^37,60^. However, the lack of a similar atlas for developing brains has hindered the systematic examination of different cell types and their evolution across neurodevelopment. Although emerging atlas frameworks have become available for the developmental mouse brain ^35,36,61^, anatomical labels with different ontologies and sparse developmental time points create significant challenges in consistently interpreting cell type specific signals, especially since the developing mouse brain rapidly evolves in structure. Our epDevAtlas resolves this by offering morphologically and intensity-averaged symmetric templates at six key ages during the critical early postnatal period, ranging from P4 to P14. Furthermore, the epDevAtlas includes 3D anatomical annotations derived from the Allen CCFv3, validated and refined using cell type-specific transgenic animals. Hence, this fills the critical need to systematically study cell type changes in early postnatal development.

Leveraging new atlases and mapping pipelines, we present detailed growth charts for GABAergic neurons and microglia, accounting for regionally distinct volumetric expansion of the brain. We found that rapid changes in volume and cell type density, including GABAergic neurons and microglia, stabilize around P12 during the first two postnatal weeks. Earlier research has demonstrated that cortical GABAergic interneurons undergo activity-dependent cell death during early postnatal periods to reach stable densities for mature inhibitory circuits ^10–12,14^ and have up to two-fold differences in their density across different areas in adult brains ^27,45^. We identify that this overall regional density difference is established as early as P4 and the density decreases approximately two-fold, reaching stability at P12. Simultaneously, microglial density in the isocortex increases about two-fold to facilitate the clearance of apoptotic neurons and promote circuit maturation ^62^. Additionally, we found that GABAergic interneuron subtypes from different developmental origins can exhibit varying magnitudes of cell density changes ^10,12^.

MGE-derived Sst neurons showed more than a two-fold reduction in select lateral association areas, while cell density reduction occurred at a much smaller magnitude in the SSp, suggesting regional heterogeneity in programmed cell death. On the contrary, CGE-derived Vip neurons established stable distribution patterns as early as P4, with minimal density changes until P14. A previous study showed that Vip neurons do not undergo activity-dependent apoptosis ^12^. This evidence suggests that Sst neurons are more plastic, in that they establish their mature density based on external stimuli, compared to Vip neurons ^10,12,63,64^. Microglia rapidly increase their cell density in the isocortex, peaking at P12, with the most substantial density change occurring in sensory cortices, while association areas showed a relatively smaller increase. This suggests that microglia play an active role in shaping activity-dependent cortical development based on external stimuli during the first two postnatal weeks ^47,56,62^. The coordinated and selective increase in microglial density within the ventrolateral area of the dorsal striatum, which receives major somatosensory cortical projections, further corroborates the notion that microglia may increase its density in sensory processing areas to facilitate activity-dependent brain development. Previous studies suggested that microglia participate in cortical processing as a negative feedback mechanism like inhibitory neurons ^65,66^. Our data raise an interesting possibility that region-specific increase of microglia density can act to regulate the influx of sensory signals from the thalamus to prevent over-excitation of the cortical circuit.

Beyond the isocortex, our results provide a comprehensive resource for examining quantitative cell type changes in other brain regions across time. Unlike cortical areas, including the hippocampus and olfactory cortices, the olfactory bulb (OB) and the striatum demonstrate continuously increasing density of Gad2 neurons, partially due to ongoing neurogenesis during the early postnatal period ^9,67,68^. Moreover, the majority of neurogenesis in the striatum is complete by birth, indicating that the rapid increase in striatal Gad2 neuronal density represents a delayed onset of Gad2 expression in medium spiny neurons, a major neuronal subtype with long-range projections ^44^. Notably, we also observed the emergence of Gad2 expression in striosomes, followed by its increase in surrounding matrix compartments. This finding aligns with the early development of striosomes during embryonic development ^43,44^. In contrast, we found an approximately two-fold decrease of Sst neurons, as one of the main interneuron subtypes in the dorsal striatum (**Extended Data Table 4**) ^44^, suggesting that the interneuron population in both cortical and striatal regions undergo significant density reductions during the early postnatal period.

Furthermore, we identified spatiotemporal patterns of specific clusters of white matter-associated microglia (WAMs) in a part of the corpus callosum and the cerebellar white matter ^69^. These microglia interact with neuronal, glial, and vascular cell types to orchestrate healthy brain development ^70–74^. Previous studies showed that WAMs might play key roles in shaping the development of white matter and survival of long-range projecting cortical excitatory neurons ^49,50,69,75,76^. Rapid reductions of WAMs and microglial expansion in the gray matter of telencephalic regions at P10 suggest that microglia may have two distinct roles in shaping the development of white and gray matter in the first and second postnatal week, respectively. The effect of microglia on cells in the local environment is limited by its vicinity with relatively short cellular processes. Hence, understanding the regional density of microglia and their changes across time can provide valuable insights into the extent of microglial influence on the development of individual brain areas ^24,75^.

### Limitation of study

Our study focused on the crucial early postnatal period between P4 and P14. Prior research has indicated that glutamatergic neurons undergo programmed cell death before P4, influencing the subsequent development of GABAergic neurons ^8^. To gain a better understanding of how excitatory and inhibitory balance is established in developing brains, future studies should explore earlier time points with glutamatergic cell types. Additionally, while our chosen Cx3cr1 mice offer relatively specific labeling of microglia, a minor population of the *Cx3cr1* gene is also expressed in other immune cells in the brain, such as border-associated macrophages ^77,78^.

Utilizing more specific transgenic reporters or combining them can greatly enhance the identification of microglia and their changes in developing mouse brains ^74,79^. Further studies are also necessary to unravel the dynamic states, functions, and implications of microglia in the developing brain and their association with related diseases ^80^.

In summary, our growth charts represent a significant stride in comprehending crucial changes in cell types that are essential for typical brain development. This resource offers a systematic framework for evaluating pathological deviations across diverse neurodevelopmental disorders. Looking ahead, we envision employing epDevAtlas to include additional cell types, such as astrocytes, oligodendrocytes, and vascular cells in further investigations. This endeavor would produce more comprehensive and nuanced brain cell type growth charts, facilitating a deeper understanding of neurodevelopment.

## Methods

### Animals

At Pennsylvania State University College of Medicine (PSUCOM), all experiments and techniques involving live animals conform to the regulatory standards set by the Institutional Animal Care and Use Committee (IACUC) at PSUCOM. For labeling GABAergic cell types during early postnatal development (P4, P6, P8, P10, P12, P14), we crossed Gad2-IRES-Cre mice (JAX, stock 028867), Sst-IRES-Cre mice (JAX, stock 013044), or Vip-IRES-Cre mice (JAX, stock 031628) with Ai14 mice, which express a Cre-dependent tdTomato fluorescent reporter (JAX, stock 007908). Heterozygous Cx3cr1-eGFP(+/–) offspring for microglia analysis were produced by crossing homozygous Cx3cr1-eGFP mice (JAX, stock 005582) with C57Bl/6J mice (JAX, stock 000664). These four animal lines were maintained and collected at PSUCOM.

Likewise, at the Allen Institute for Brain Science (referred to as the ‘Allen Institute’), all animal experiments and techniques have been approved and conform to the regulatory standards set by the Institutional Animal Care and Use Committee (IACUC) at the Allen Institute. For labeling cortical layer cell types at P6, P10, and P14, we used nine mouse genotypes. Slc32a1-IRES-Cre mice (JAX, stock 016962) were crossed with Ai65 reporter mice (JAX, stock 021875) and further crossed with Lamp5-P2A-FlpO mice (JAX, stock 037340) to produce triple transgenic offspring for layer 1 (L1) Slc32a1+/Lamp5+ cells. Layer 2/3 (L2/3) Calb2+ cells were labeled by crossing Calb2-IRES-Cre mice (JAX, stock 010774) with Ai14 reporter mice. Layer 4 (L4) Nr5a1+ cells were labeled by crossing Nr5a1-Cre mice (Mutant Mouse Resource & Research Center, stock 036471-UCD) with Ai14 reporter mice. For layer 5 (L5) Rbp4+ cells, Rbp4-Cre KL100 mice (Mutant Mouse Resource & Research Center, stock 037128-UCD) were crossed with Ai14, Ai193 (JAX, stock 034111), or Ai224 reporter mice (JAX, stock 037382). Layer 6 (L6) Ntsr1+ cells were labeled by crossing Ntsr1-Cre GN220 mice (Mutant Mouse Resource & Research Center, stock 030648-UCD) with Ai14 reporter mice. Layer 6b (L6b) Cplx3+ cells were labeled by crossing Cplx3-P2A-FlpO mice (JAX, stock 037338) with Ai193 or Ai227 (JAX, stock 037383) reporter mice.

Genotyping was performed by PCR of tail biopsy genomic DNA for certain mouse lines. For mice younger than P6, Rbm31-based genotyping was used since visual identification of neonatal mouse sex based on anogenital distance is challenging ^81^. Detailed information on transgenic reporter lines and animal numbers is available in Extended Data Table 1. All mice had access to food and water ad libitum, were maintained at 22–25 °C with a 12-hour light/12-hour dark cycle, and both male and female mice were included in the study, with each animal used once for data generation.

### Brain collection, embedding, STPT imaging, and 3D reconstruction

The collection and STPT imaging of mouse brains have been extensively detailed in our protocol paper ^82^. Briefly, animals were deeply anesthetized with a ketamine and xylazine mixture (100 mg/kg ketamine, 10 mg/kg xylazine, intraperitoneal injection) before perfusion. Transcardiac perfusion involved washing out blood with isotonic saline solution (0.9% NaCl) followed by tissue fixation with freshly made 4% PFA in phosphate buffer (0.1 M PB, pH 7.4). Post-fixation occurred by decapitating the heads and storing them in 4% PFA for 2 days at 4°C. This was followed by careful brain dissection to ensure preservation of all structures. The brains were then stored in 0.05 M PB (pH 7.4) until STPT imaging preparation. Animals with incomplete perfusion or dissection were excluded from imaging and analysis.

At PSUCOM, precise STPT vibratome cutting was achieved by embedding fixed brains in 4% oxidized agarose in custom-built molds, ensuring consistent 3D orientation ^82^. Cross-linking was achieved by incubating samples in 0.05 M sodium borohydride solution at 4°C overnight before imaging. STPT imaging was performed using a TissueCyte 1000 (TissueVision) with a 910 nm two-photon laser excitation source (Chameleon Ultra II, Coherent). Green and red signals were simultaneously collected using a 560 nm dichroic mirror. Sampling rate (pixel size) was 1 × 1 μm (xy) and image acquisition occurred at intervals of 50 μm (z). We utilized custom-built algorithms to reconstruct STPT images into 3D volumes ^45,82^.

At the Allen Institute, fixed brain samples were embedded in 4% oxidized agarose and incubated overnight in acrylamide solution at 4°C before heat-activated polymerization the following day (detailed protocol available at https://www.protocols.io/view/tissuecyte-specimen-embedding-acrylamide-coembeddi-8epv512njl1b/v4). Embedded samples were stored in 50 mM PB prior to STPT imaging using a TissueCyte 1000 with a 925 nm two-photon laser excitation source (Mai Tai DeepSee, Spectra-Physics). Green and red signals were simultaneously collected using a 560 nm dichroic mirror. Sampling rate (pixel size) was 0.875 μm × 0.875 μm (xy) and image acquisition occurred at intervals of 50 μm (z). Acquired images were transferred to the PSUCOM for further analysis.

### epDevAtlas Template Generation

Background channels of STPT-imaged data were used to construct morphologically averaged symmetric reference brain templates at ages P4, P6, P8, P10, P12, and P14. Templates were primarily generated using Vip-Cre;Ai14 mouse brains as the tdTomato signal from the fluorescent reporter is minimally visible once resampled to the template resolution of 20 μm × 20 μm × 50 μm (XYZ in the coronal plane). The P6 template was supplemented with data from Gad2-Cre;Ai14 and Sst-Cre;Ai14 data. The sample size per template varied between 6 to 14 from males and females.

To obtain symmetric templates, each preprocessed image underwent duplication and reflection across the sagittal midline. This step effectively doubled the number of input datasets used in the template construction pipeline, ensuring bilateral congruence. Applied Normalization Tools (ANTs) was utilized for registration-based methods to create a morphologically averaged symmetric template for each developmental age ^83,84^. Morphologically averaged symmetric templates were created on Penn State’s High-Performance Computing system (HPC) for each developmental age guided by the ANTs function, ’antsMultivariateTemplateConstruction2.sh’ as described in Kronman et al ^36^. The procedure started by creating an initial template estimate from the average of input datasets. Following initialization, (1) Each input image was non-linearly registered to the current template estimate. (2) The non-linearly registered images were voxel-wise averaged. (3) The average transformation derived from registration was applied to the voxel-wise average image generated in the previous step, thereby updating the morphology of the current template estimate. The iterative process continued until the template’s shape and intensity values reached a point of stability.

### Down Registration of CCFv3 Labels to Templates

The P56 STPT Allen CCFv3 anatomical labels (RRID:SCR_020999) were iteratively down registered to each developmental timepoint represented by our STPT templates using manually drawn landmark registration ^36^. The CCFv3 template was initially registered to the P14 STPT template utilizing non-linear methods. Subsequently, registration quality was assessed by superimposing the warped CCFv3 onto the P14 STPT template in ITK-SNAP ^85^ to identify critical misaligned landmark brain regions visually. Misaligned regions and whole brain masks were segmented for both templates using Avizo (Thermo Fisher Scientific), a 3D image visualization and analysis software. The segmented regions were then subtracted from the brain masks, creating modified brain masks with boundaries defined around the misaligned brain regions. Next, we performed linear registration of the CCFv3 and P14 STPT modified brain masks, followed by equally weighted non-linear registration of the template images and their corresponding modified brain masks. Finally, we applied the transformation derived from landmark registration to the CCFv3 anatomical labels, moving them to the P14 STPT template morphology. This process was repeated sequentially to align the anatomical labels from the P14 STPT template to the P12 STPT template, then to the P10 STPT template, and so forth until the P4 STPT template morphology was reached.

### Cell detection, image registration, and 3D cell counting

We developed a flexible and automatic workflow with minimal annotations and algorithmic training based on our previous cell density mapping methods ^27,35,45^. We employed ilastik as a versatile machine learning tool using random forest classification for signal detection ^86^, instead of more resource-intensive deep learning approaches that necessitate larger training sets and increased computational resources. Integrating ilastik into the automatic workflow, we designed our algorithms to perform parallel computations to detect each pixel with the maximum likelihood of it belonging to a cell, brain tissue, or empty space ^29^. Detected signals that were deemed too small for cells were considered artifacts and discarded. Finally, we recorded the location of the center of mass (centroid) for each cell cluster. We performed image registration to map cell detection results to an age-matched epDevAtlas template using elastix ^87^. The number of centroids was calculated for each brain region to generate 2D cell counting, which then was converted to 3D cell counting pre-established conversion factors (1.4 for cytoplasmic signals and 1.5 for nuclear signals) ^27^. To calculate the anatomical volume from each sample, the epDevAtlas was first registered to individual samples using elastix and anatomical labels were transformed based on the registration parameters. Then, the number of voxels associated with specific anatomical IDs was used to estimate the 3D volume of each anatomical area. 3D cell counting per anatomical regional volume (mm^3^) was used to calculate the density.

### Data visualization, including isocortical flatmap

To visualize cell type density across different isocortical regions, we utilized a custom MATLAB script to map Allen CCFv3 registered signals onto a 2D projected isocortical flatmap ^45^. First, an isocortical flatmap was generated for individual sample datasets, using 3D counted cell data registered to the Allen CCFv3. For each postnatal timepoint per cell type, the flatmap images were averaged using a MATLAB script. Then, the averaged flatmaps were normalized for isocortical volume since the data were registered to Allen CCFv3 adult brain template. For image normalization, multiplication factors were determined for each postnatal timepoint (P4 to P14) by taking the average isocortical volume from age-matched developing brains and dividing those values by the average adult mouse isocortical volume. After normalization, each isocortical flatmap represented the average density of each cell type at a specific postnatal timepoint. To visualize cell type density across major regions of the entire brain, we created cell density maps using the bubble chart template in Excel (Microsoft, v.16.72). The size of each bubble corresponds to cell density values. Additionally, we plotted region-specific cell densities over time for each cell type using Prism (GraphPad, v.9.5.1).

### Statistical analyses

All data were presented as the mean ± standard deviation (SD). Significance was determined by a p-value of less than 0.05. Prior to performing statistical analyses (GraphPad Prism, v.9.5.1), the datasets were assessed for normality and homogeneity of variance to check if the assumptions for parametric tests were met. Since the datasets did not meet these criteria, differences between groups were analyzed using Welch’s one-way ANOVA followed by a non-parametric Dunnett’s post hoc test. Adjusted p-values from the multiple comparisons test were used to determine significance. Quantified cell type density data were collected, organized, and presented in Extended Data Tables 3, 4, 5, and 6 (Microsoft Excel, v.16.72).

To compare developmental variation in isocortical cell densities of Gad2, Vip and Sst neurons and Cx3cr1 microglia across anatomical space, we performed one-way functional ANOVA using the scikit-fda package, which implements functional data analysis (FDA) in the Python scikit-learn machine learning framework ^88^. The FDA evaluates each observation as a function of a variable. One-way functional ANOVA calculates the sample statistic Vn, which measures the variability between groups of n samples, and implements an asymptotic method to test the null hypothesis that the Vn is equivalent to an asymptotic statistic V, where each sample is replaced by a gaussian process, with mean zero and the original covariance function. The sampling distribution of the asymptotic statistic V is created by repetitive simulation using a bootstrap procedure. For post hoc analysis of ANOVA results where the null hypothesis was rejected, i.e. p < 0.05, we employed pairwise permutation t-tests for all age groups, which create a null distribution for a test of no difference between pairs of functional data objects, using a MATLAB implementation of FDA.

### Data and code availability

The epDevAtlas is an openly accessible resource package including age-matched templates and anatomical labels which can be viewed and downloaded via https://kimlab.io/brain-map/epDevAtlas/

Full resolution and mapped cell type data can be also found at https://kimlab.io/brain-map/epDevAtlas/.

All available data and code will be deposited in public data repository (e.g., Mendeley data, GitHub) upon publication.

## Acknowledgments

This work was supported by a grant from the National Institute of Mental Health (RF1MH12460501, R01NS108407). Our deepest thanks to members of the BRAIN Initiative Cell Census Network for their insights. We express gratitude to all members of the Yongsoo Kim Lab for their motivation, commitment, and knowledge. We are utmost grateful toward all team members from the Allen Institute for Brain Science for graciously providing their time and efforts, including but not limited to animal husbandry, microscopy imaging (Nhan-Kiet Ngo and Nadezhda I. Dotson), and data management. We thank Julie Nyhus from the Allen Institute for managing all collaborative efforts during this project. Additionally, we thank BioRender.com for their illustration generation platform and the high-performance computing (HPC) center at PSUCOM.

## Contributions

Corresponding author Y.K. conceived the project, supervised data generation and analysis, and edited manuscript. J.K.L. collected data, performed data analysis, and wrote the manuscript with help from all authors. J.A.M., Y.B., M.T., S.W., H.Z., B.T., and L.N. provided experimental support, generated data, and performed quality control. D.P. assisted in conducting statistical analyses. F.K. helped to create epDevAtlas and Y.T. developed the cell counting pipeline. D.J.V. generated the web visualization platform.

## Extended Data Figures

**Extended Data Figure 1.**
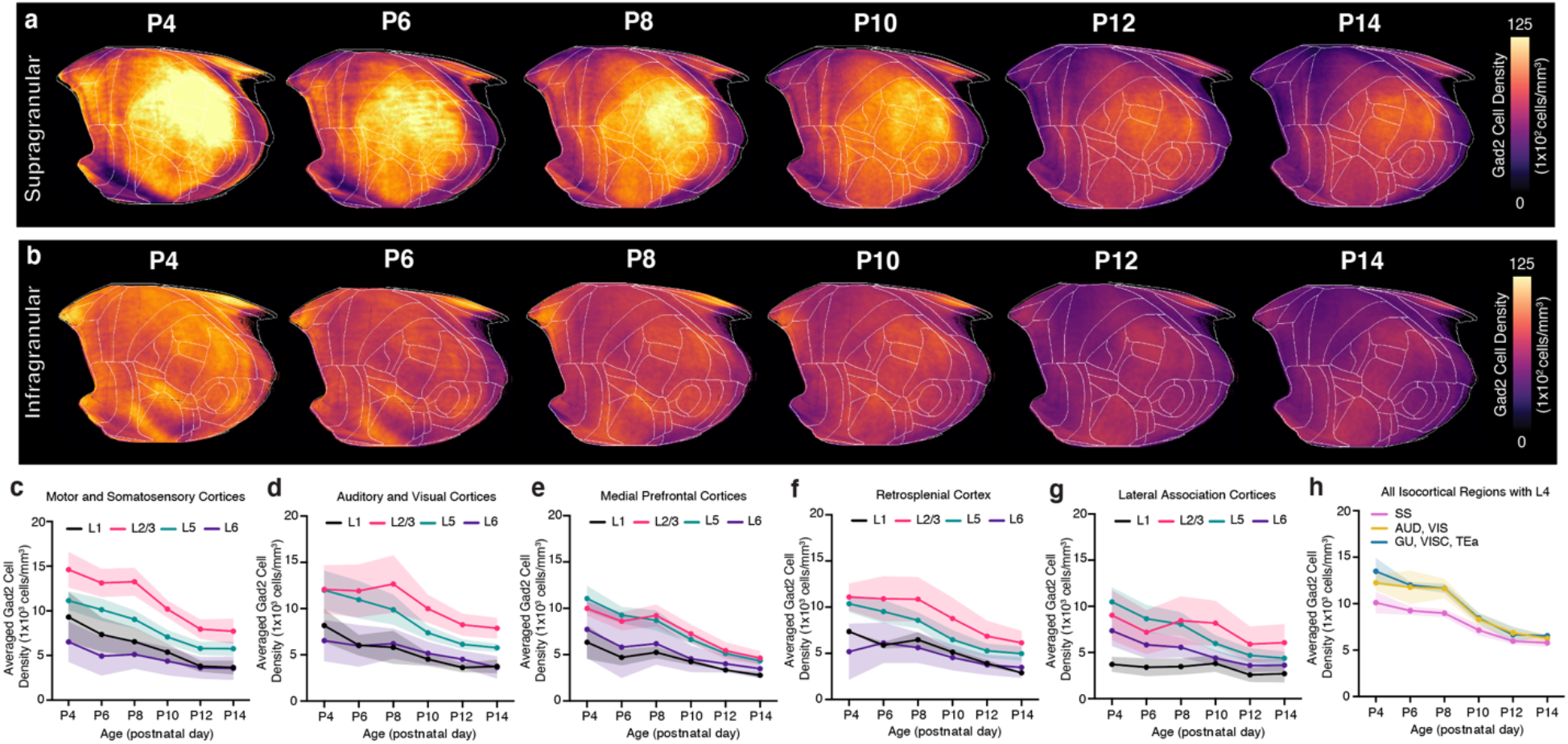
Cortical layer developmental mapping of Gad2 neurons. **a-b**, Isocortical flatmaps of Gad2 neuronal densities in **(a)** supragranular (L1, L2/3, L4) and **(b)** infragranular (L5, L6) cortical layers. **c-g,** Layer-specific trajectories of Gad2 cell density in L1, L2/3, L5, and L6 of the isocortex divided into regional subgroups based on their functional and anatomical connectivity: **(c)** motor and somatosensory, **(d)** auditory and visual,€) medial prefrontal, **(f)** retrosplenial, and **(g)** lateral association areas. **h,** Gad2 cell density in isocortical regions containing L4, which includes somatosensory (SS), auditory (AUD), visual (VIS), gustatory (GU), visceral (VISC), and temporal association (TEa) areas. Data are reported as mean ± s.d. (shaded area between error bars). See Extended Data Table 3 for Gad2 cell counts, density, and volume measurements.

**Extended Data Figure 2.**
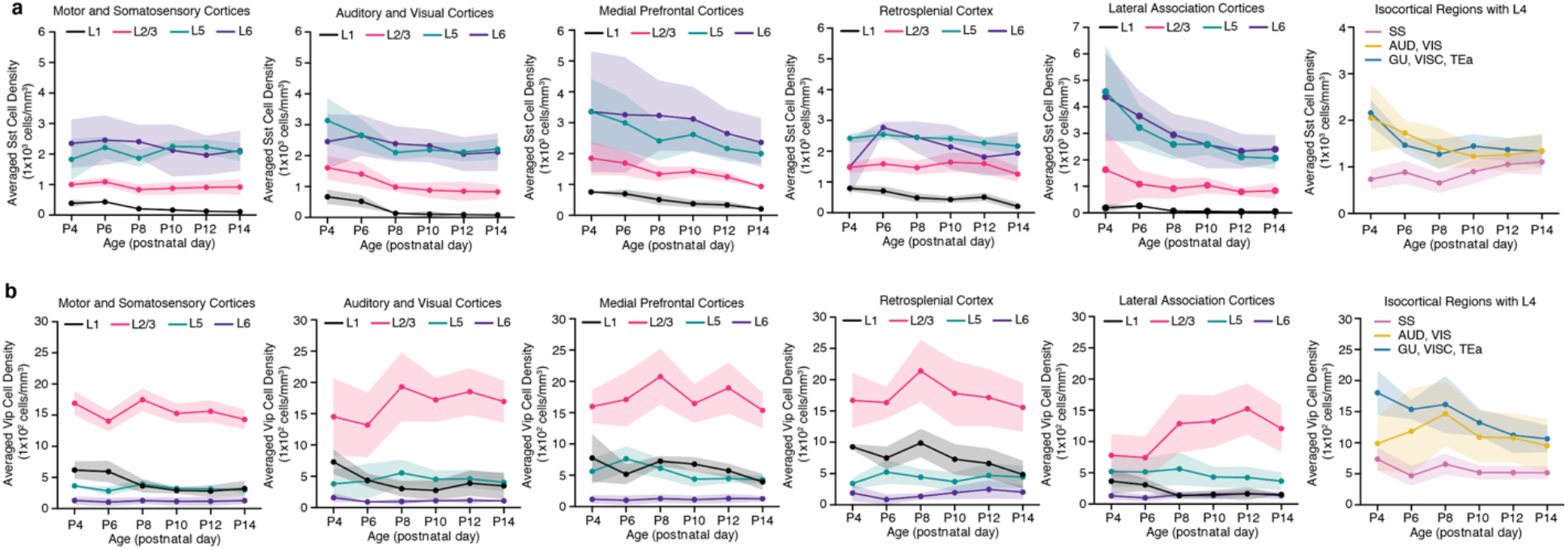
Cortical layer developmental mapping of Sst and Vip interneurons. **a-b**, Layer-specific trajectories of (**a**) Sst cell density and (**b**) Vip cell density in the isocortical areas. Data are reported as mean ± s.d. (shaded area between error bars). See Extended Data Tables 4 and 5 for cell counts, density, and volume measurements for Sst and Vip interneurons, respectively.

**Extended Data Figure 3.**
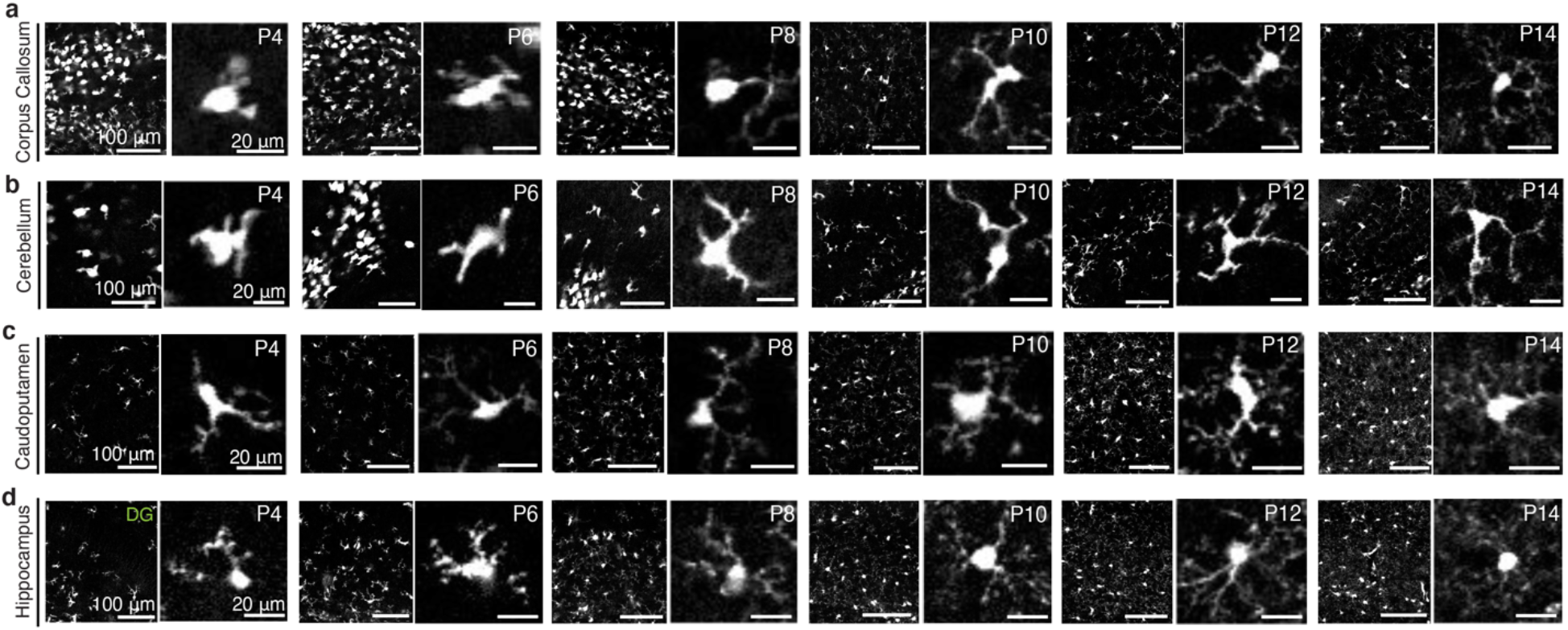
Spatiotemporal changes in early postnatal microglial morphology. **a-d**, Representative STPT images of Cx3cr1 microglia in various brain regions at P4, 6, 8, 10, and 14. Low (left) and high magnification (right) images from each age. (**a**) Amoeboid-shaped, white matter tract-associated microglia (WAMs) with large somas and short, thick branches are present in the corpus callosum from P4 until P8, before adopting a more ramified morphology with longer, extended processes at P10 and onward. (**b**) Likewise, these WAMs with similar morphological specifications outlined in (**a**) are present in the cerebellar white matter. Microglia in gray matter brain regions, such as the (**c**) caudoputamen and the (**d**) dentate gyrus (DG) of the hippocampus exhibit different morphological changes compared to WAMs, with smaller somas and short processes that become increasingly larger and longer, respectively.

**Extended Data Figure 4.**
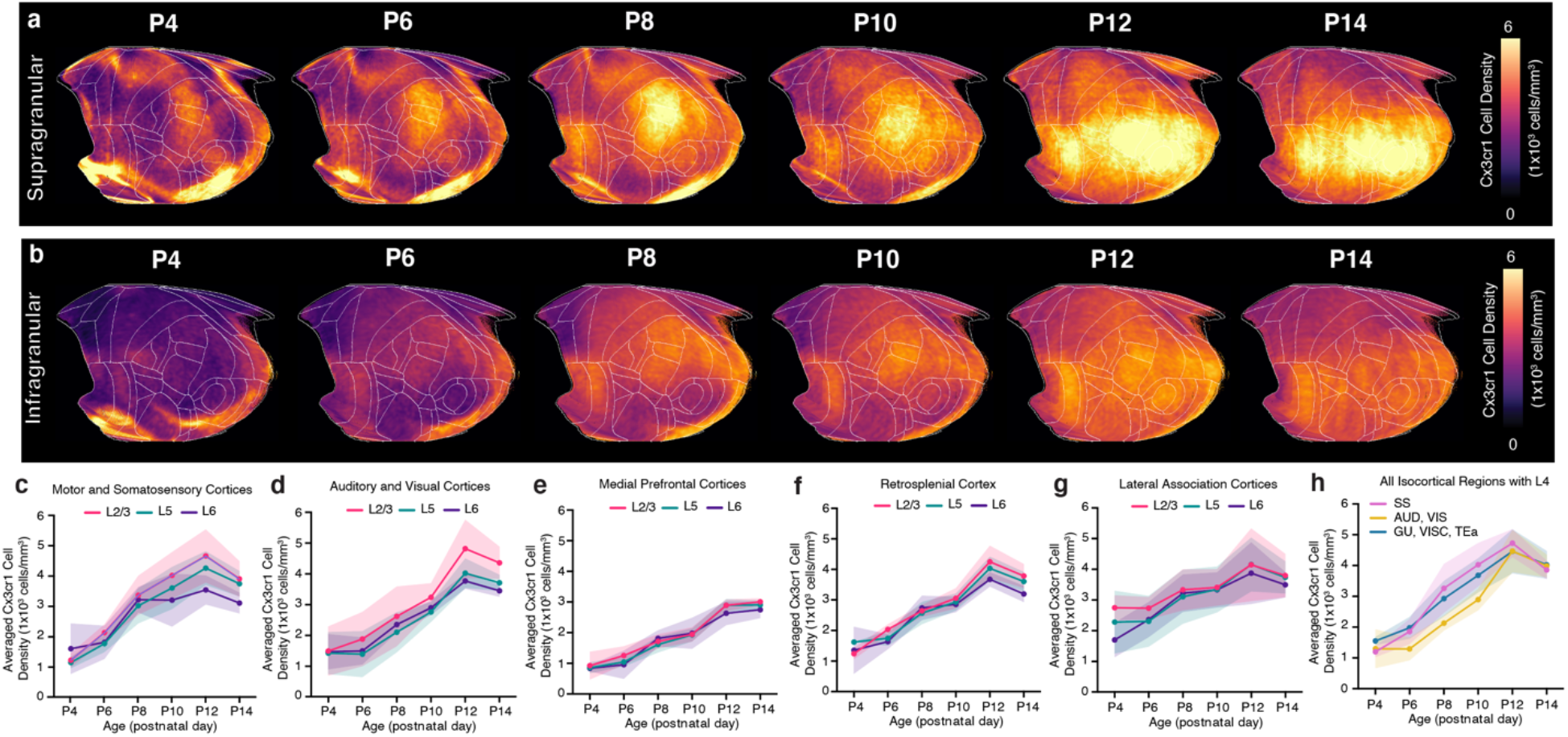
Cortical layer developmental mapping of microglia. **a-b**, Isocortical flatmaps of Cx3cr1 microglial densities ranging from P4 to P14, showing the distinct spatial distribution patterns between **(a)** supragranular (L1, L2/3, L4) and **(b)** infragranular (L5, L6) cortical layers. **c-g,** Layer-specific trajectories of averaged Cx3cr1 microglial density in L1, L2/3, L5, and L6 of the isocortex divided into regional subgroups based on their functional and anatomical connectivity: (**c**) motor and somatosensory, (**d**) auditory and visual, (**e**) medial prefrontal, (**f**) retrosplenial, (**g**) lateral association areas, **(h)** isocortical regions containing L4. Data are reported as mean ± s.d. (shaded area between error bars). See Extended Data Table 6 for Cx3cr1 microglia counts, density, and volume measurements.

